# Enhancing Drug-Drug Interaction Prediction Using Deep Attention Neural Networks

**DOI:** 10.1101/2021.03.16.435553

**Authors:** Shichao Liu, Yang Zhang, Yuxin Cui, Yang Qiu, Yifan Deng, Wen Zhang, Zhongfei Zhang

**Affiliations:** College of Informatics, Huazhong Agricultural University, Wuhan 430070; Electronic Information School, Wuhan University, Wuhan 430072; Computer Science Department, Binghamton University, Binghamton, NY, USA

**Keywords:** drug-drug interactions, deep attention neural networks

## Abstract

Drug-drug interactions are one of the main concerns in drug discovery. Accurate prediction of drug-drug interactions plays a key role in increasing the efficiency of drug research and safety when multiple drugs are c o-prescribed. With various data sources that describe the relationships and properties between drugs, the comprehensive approach that integrates multiple data sources would be considerably effective in making high-accuracy prediction. In this paper, we propose a Deep Attention Neural Network based Drug-Drug Interaction prediction framework, abbreviated as DANN-DDI, to predict unobserved drug-drug interactions. First, we construct multiple drug feature networks and learn drug representations from these networks using the graph embedding method; then, we concatenate the learned drug embeddings and design an attention neural network to learn representations of drug-drug pairs; finally, we adopt a deep neural network to accurately predict drug-drug interactions. The experimental results demonstrate that our model DANN-DDI has improved prediction performance compared with state-of-the-art methods. Moreover, the proposed model can predict novel drug-drug interactions and drug-drug interaction-associated events.

## 1 Introduction

Drug-drug Interactions (DDIs) may occur when multiple drugs are co-prescribed. Although DDIs may have beneficial effects, they sometimes have serious adverse effects and lead to drug withdrawal from the market [1]. Drug-drug interaction (DDI) prediction can help reduce the probability of adverse reactions and optimize the drug development and post-marketing surveillance process.

Clinical trials are time-consuming, expensive, and infeasible when facing the large-scale data and limitations of experimental conditions. Therefore, the researchers introduce a lot of computational methods to accelerate the prediction process. The existing computational DDI prediction methods can be roughly divided into five categories: literature extraction-based, matrix factorization-based, similarity-based, network-based and deep learning-based methods.

Literature extraction-based methods take the extraction of DDIs as a multi-class classification task, which generally extract information from informative sentences of literature, then detect candidate DDIs and classify them. Chowdhury et al. [2] proposed a multi-phase relation extraction approach, which used hybrid kernel-based support vector machine to extract the possible relationships between drug-drug pairs. Sun et al. [3] constructed a deep architecture with multiple layers of small convolutions based on a convolutional neural network to extract features automatically. Kim et al. [4] utilized a linear kernel-based classifier and added lexical and syntactic features to improve the accuracy of prediction. Zhang et al. [5, 6] proposed a hierarchical RNNs-based model, which combined short dependency path (SDP) and sentence sequence to obtain semantic information and used attention mechanism to map representations. Jiang et al. [7] introduced a skeleton LSTM method to effectively extract the internal structure of DDIs.

Matrix factorization-based methods decompose the drug-drug interaction matrix into several matrices, which reconstruct the interaction matrix to predict DDIs. Zhang et al. [8] introduced the manifold regularization based on drug features and proposed a novel matrix factorization method named MRMF to predict potential DDIs. Shi et al. [9] adopted the triple matrix factorization model to predict not only binary DDIs but comprehensive DDIs (i.e., enhancive and degressive DDIs) further. Yu et al. [10] designed DDINMF based on semi-nonnegative matrix factorization which decomposes the DDI adjacent matrix through the regular nonnegative matrix factorization (NMF) for enhancive and degressive DDI prediction.

Similarity-based methods assume that similar drugs could interact with the same drug. Gottlieb et al. [11] constructed a model named INDI which calculated seven types of similarity and used a weighted logistic regression classifier to predict drug-drug interactions. Cheng et al. [12] integrated a variety of drug-drug similarities to describe drugdrug pairs, and used five classifiers to build the prediction models. Foukoue et al. [13] proposed a framework named Tiresias which constructed a knowledge graph through semantic integration of data and utilized the graph to calculate similarities among drugs.

Network-based methods consider the high-order similarity of drugs and the propagation of similarity, or infer from the network structure directly. Zhang et al. [14] introduced a label propagation framework which computed the similarities of side effects, off label side effects and chemical structure, and paid attention to high-order similarity. Park et al. [15] used a random walk method to simulate signaling propagation on the protein-protein interaction (PPI) network. Lee et al. [16] constructed a heterogeneous bioinformatics network and adopted graph traversal algorithms to find paths of drug-drug pairs. Huang et al. [17] proposed a metric named S-score to measure the strength of network connections and obtained accurate prediction results through the Bayesian probability model.

Moreover, deep learning-based methods, which are specialized in extracting high-quality drug features, have been widely applied in the field of biology [18, 19] and have achieved promising results. Karim et al. [20] constructed a knowledge graph which considered combining CNN and LSTM model to extract local and global features of drugs through the network. Chu et al. [21] utilized a factorization auto-encoder to learn the representations of complex nonlinear drug interactions. Liu et al. [22] introduced the multimodal deep auto-encoder named DDI-MDAE based on shared latent representation to predict DDIs.

Many existing methods are proposed to predict whether drugs interact or not. In fact, diversified sub-types of interactions could instruct further research on interaction events [23, 24]. Hence, DDI-associated events prediction has aroused some attention. Ryu et al. [25] proposed a deep learning method using structural information and a deep neural network to predict 86 important DDI types. Lee et al. [26] used auto-encoders to reduce dimensions of similarity files and applied a deep neural network to predict 106 types of DDIs. Deng et al. [27] proposed a multimodal deep learning framework which utilized multiple drug features to predict 65 categories of DDI events.

Recent studies have highlighted the importance of integrating heterogeneous drug features for the prediction of DDIs [28, 29]. However, integrating multiple diverse drug features in drug-drug interaction prediction is a challenging task. Heterogeneous drug features could be correlated and have redundant information, which may affect the performances of conventional classifiers. Hence, an effective framework of integrating heterogeneous features is in demand. Deep Neural Network (DNN) is one of the representative algorithms of deep learning, which aims to use multiple nonlinear and complex processing layers to model high-level drug features. Thus, it is a suitable model for learning high-level representations from multiple drug features.

In this paper, we propose a Deep Attention Neural Network based DDI prediction framework, abbreviated as DANN-DDI. Firstly, drug features are formulated as drug feature networks, and the graph representation learning method SDNE [30] is utilized to learn drug embeddings from these networks. Secondly, the drug embeddings are concatenated, and the attention neural network, which takes into account different contributions of different features and their dimensions, is designed to learn the representations of drug-drug pairs. Finally, the representations of drug-drug pairs are fed into a deep neural network to predict potential drug-drug interactions. The experimental results demonstrate that DANN-DDI could combine diverse drug information attentively, deliver high-accuracy performances, and outperform benchmark methods.

In summary, the main contributions of DANN-DDI are as follows: (1) We introduce a deep attention neural network framework for drug-drug interaction prediction, which can effectively integrate multiple drug features; (2) We employ the attention neural network to learn attention vectors of each of the specific drug-drug pairs; (3) Experimental results demonstrate that DANN-DDI can predict novel drug-drug interactions and DDI-associated events.

## 2 Materials and methods

### 2.1 Dataset Description

DrugBank [31] database integrates bioinformatics and chemical informatics resources to provide detailed drug data, including drug chemical substructures, targets, enzymes, pathways and drug-drug interactions. In this paper, we adopt the drug dataset with multiple drug features obtained from the DrugBank database released in April 2018 (version 5.1.0). Then, we use the ID mapping server to map the target proteins of the drugs into the KEGG drug database [32] for obtaining drug pathways. Our dataset contains 841 drugs with 619 chemical substructures, 1333 targets, 214 enzymes and 307 pathways, which is described in Table 1. The 841 drugs have 353,220 drug-drug pairs, including 82,620 known pairwise DDIs.

**TABLE 1.** The description about drug features and sources

| Data type | Database | Reference | Description |
| --- | --- | --- | --- |
| Drug | DrugBank | [31] | 841 drug types |
| Target | DrugBank | [31] | 1333 target types |
| Enzyme | DrugBank | [31] | 214 enzyme type |
| Pathway | KEGG | [32] | 307 pathway types |
| Substructure | PubChem | [33] | 619 substructure types |

### 2.2 Overview of Methods

In this paper, we propose a deep attention neural network framework named DANN-DDI to predict potential interactions between drug-drug pairs in multiple drug feature networks.

As described in Figure 1, DANN-DDI consists of three components, i.e. drug feature learning component, drugdrug pair feature learning component and drug-drug interaction prediction component. The drug feature learning component first constructs drug feature networks, and then learn the representations of drugs from these networks using graph representation learning (Figure 1a). The drug-drug pair feature learning component concatenates the five embeddings to obtain the comprehensive embeddings for drugs and designs an attention neural network to learn drug-drug pairs’ representations (Figure 1b). The drug-drug interaction prediction component uses a deep neural network to predict potential drug-drug interactions, which uses the representations of drug-drug pairs as input (Figure 1c).

**Fig. 1.**
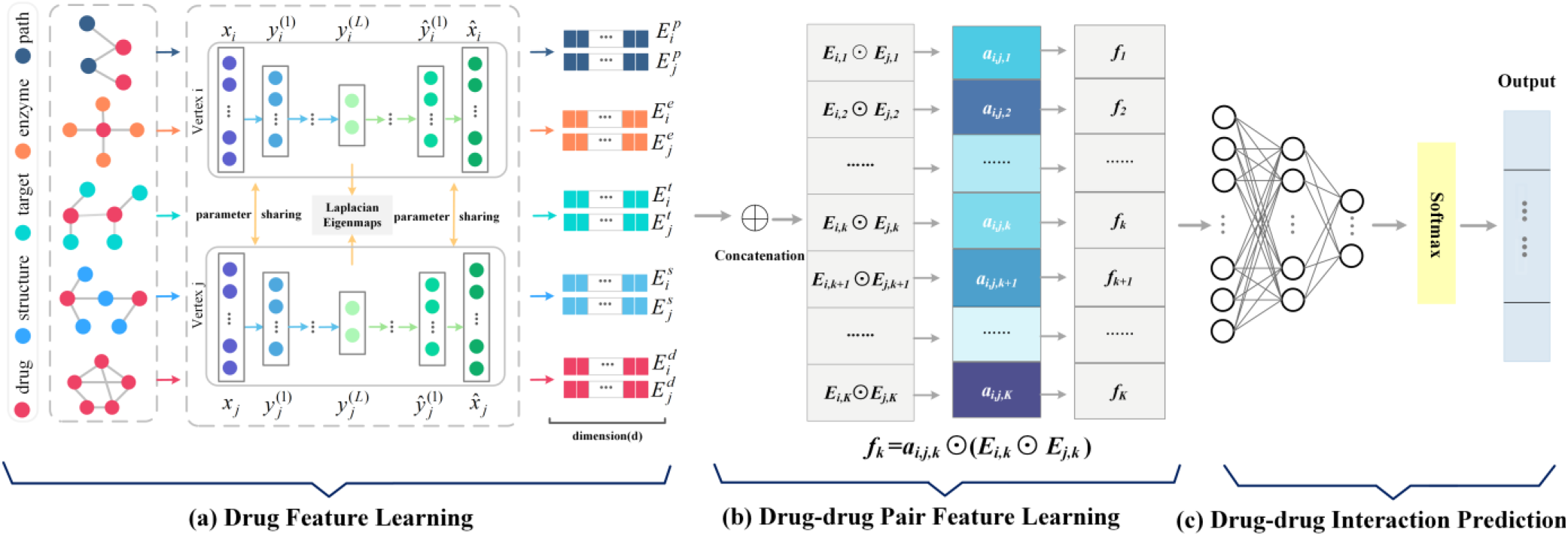
An overview of DANN-DDI. (a) Drug Feature Learning. (b) Drug-drug pair feature learning. (c) Drug-drug interaction prediction.

### 2.3 Drug Feature Learning

In this work, we collect five drug features, i.e. chemical substructures, targets, enzymes, pathways, and drug-drug interactions. The basic idea of the drug feature learning is described in Figure 1a.

#### 2.3.1 Construction of Drug Feature

Due to the different effects of drug features on the performances of DDI prediction, we consider all these drug features to build five drug feature networks: drug-substructure network, drug-target network, drug-enzyme network, drug-pathway network and drug-drug interaction network. Each drug feature network includes drug nodes, feature nodes, and the links between them. The links of the networks represent associations between drugs and features. For example, a drug could have some substructures, and thus a drug node is linked to some substructure nodes in the drug-substructure network.

#### 2.3.2 Graph Representation Learning

The graph embedding or representation learning is to learn the feature vectors of nodes in networks, and gains more and more attentions in bioinformatics [34]. Many graph representation learning methods have been proposed, and studies have shown that the structural deep network embedding (SDNE) could achieve competitive performances. Therefore, we adopt SDNE to learn the embeddings of drug nodes from drug feature networks.

SDNE can preserve the first-order and second-order proximity simultaneously. The first-order proximity constrains the similarity of the latent representations (i.e., 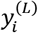 and 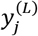) of a pair of vertexes (i.e., vertex *i* and vertex *j*). The second-order proximity is reconstructed by minimizing the reconstruction error of the output 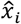 and the and the input *x_i_* More details can be found in [30].

We extract the representations of drug nodes in five drug feature networks. Suppose that we have *n* drugs; the five drug embeddings from above networks are denoted as 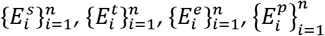 and 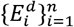 respectively.

### 2.4 Drug-drug Pair Feature Learning

Motivated by the successful applications of attention networks [6, 35–38], we introduce the drug-drug pair feature learning component in Figure 1b, which is an attention neural network to learn representations of drug-drug pairs by fusing learned representations of drugs from drug feature networks.

First, we concatenate the five embeddings 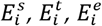,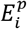 and 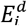, then obtain a comprehensive vector 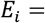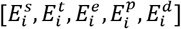 for drug *i*, *i* = 1,2,…, *n*. The comprehensive vectors combine diverse information learned from drug feature networks. Let *K* denote the dimensions of the comprehensive vectors.

Then, we learn the representations of drug-drug pairs from the comprehensive vectors of individual drugs. The representation of a drug-drug pair (drug *i* and drug *j*) is defined as follows:

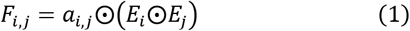

where ⨀ is the element-wise product. *a*_*i, j*_ = (*a*_*i*,*j*,1_, *a*_*i*,*j*,2_, ⋯, *a*_*i*,*j*,*K*_) is a *K*-dimensional attention vector to capture the different importance of *K* dimensions in *E*_*i*_⨀*E*_*j*_. Specifically, *F*_*i*,*j*_ = (*f*_1_, *f*_2_, ⋯, *f*_*K*_) and *f*_*k*_ = *a*_*i*,*j*,*k*_⨀(*E*_*i*,*k*_⨀*E*_*j*,*k*_). *a*_*i*,*j*,*k*_ is the attention weight responsible for the *k*th dimension, *k* = 1,2,…, *K*.

Next, we calculate the attention vector *a*_*i*,*j*_ as follows:

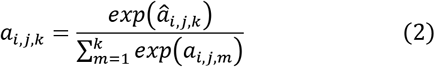

where 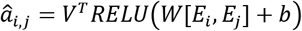 and *RELU* is an activation function [39]. *b* and *W* are the bias vector and the weight matrix, and *V*^*T*^ is the weight vector. [*E*_*i*_, *E*_*j*_] denotes the concatenation of *E*_*i*_ and *E*_*j*_. Attention vector *a*_*i*,*j*,*k*_ is used to capture the importance on dimension *k* of the representation of a drug-drug pair (drug *i* and drug *j*).

Therefore, the drug-drug pair feature learning component employs the attention neural network to learn representations of drug-drug pairs, which takes into account different contributions of different features and their dimensions. The attention neural network outputs representations of drug-drug pairs.

### 2.5 Drug-drug interaction prediction

The prediction of drug-drug interactions can be considered as a binary classification problem. The drug-drug interaction prediction component uses representations of drug-drug pairs to predict potential DDIs.

The drug-drug interaction prediction component is designed as follows. As shown in Figure 1c, the input layer of the network has the same dimensions as those of the representations of drug-drug pairs. Then, the inputs pass through multiple fully connected hidden layers. The output layer contains two neurons, which denote two classes of drug-drug pairs (interaction or no-interaction). The softmax function is used for the output nodes to generate probabilities. We adopt the Rectified Linear Unit (RELU) [39] as the activation function for all the hidden layers.

### 2.6 Model Optimization

DANN-DDI consists of three components. The drug feature learning component is trained and optimized first. Then, drug-drug pair feature learning component is trained and optimized along with the drug-drug interaction prediction component in an end-to-end way.

To optimize the DANN-DDI models, we select the binary cross-entropy loss function and utilize Adam optimizer [40] with default parameters to optimize the drug-drug interaction prediction component. Between hidden layers, batch normalization layers are adopted to accelerate the convergence, and dropout layers [41] are adopted to avoid overfitting and improve generalization ability.

## 3 Experiment

### 3.1 Evaluation Metrics

In the previous work, 5-fold cross validation was adopted to evaluate prediction models [42, 43]. In this paper, we adopt *K*-fold cross validation (i.e. 5-CV) to evaluate DDI prediction models. We randomly split the known drug-drug interactions into *K* subsets with an equal size. In each fold, one subset and all unlabeled data are selected as the testing set and the rest subsets and all unlabeled data as the training set. We use the known drug-drug interactions to train the prediction model and predict unobserved interactions between drugs.

Here, several evaluation metrics are used to measure the performances of the prediction model, i.e., the area under the precision-recall curve (AUPR), the area under ROC curve (AUC), F-measure (F), the accuracy (ACC), precision and recall. In our work, known drug-drug interactions are a small proportion of all drug-drug pairs. Therefore, we take AUPR which focuses on positive examples as the primary evaluation metric.

### 3.2 Parameter Discussion

There are four key parameters: the dimension of drug embeddings *d*, the number of DNN hidden layers *α*, the total epochs of DNN training *β* and the dropout rate *ɛ* for DNN hidden layers. Here, we discuss the influence of these parameters on the performance of DANN-DDI.

We consider the combinations of parameters: *d* ∈{32, 64, 128, 192, 256, 300}, α ∈{4, 5, 6, 7, 8}, *β* ∈{50, 100, 150, 200, 250} and *ɛ* ∈{0, 0.1, 0.2, 0.3, 0.4, 0.5}. We build the DANN-DDI models using different parameter combinations, and DANN-DDI models are evaluated using 3-CV. The AUPR scores (primary metric) and AUC scores are adopted to measure the performances of prediction models.

Among all parameter combinations, DANN-DDI achieves the best performance when *d* = 128, *α* = 7, *β* = 150 and *ɛ* = 0.4. Then, we fix three parameters and discuss the influence of the remaining parameter. When fixing *α*, *β* and *ɛ*, the AUPR scores and AUC scores of DANN-DDI model achieve the best performances when the dimension *d* of drug embeddings equals to 128 or 256 as shown in Figure 2a. When fixing *d*, *β* and *ɛ*, we set the number of neurons in the last hidden layer to 64 and consider the influence of *α* on DANN-DDI. The AUPR score of DANN-DDI increases as *α* increases, and it decreases after reaching the peak as given in Figure 2b. When *d*, *α* and *ɛ* are fixed, Figure 2c shows that the performance of DANN-DDI increases when *β* increases from 50 to 150 and decreases slightly after that, and then increases when *β* increases further. As shown in Figure 3d, when *d*, *α* and *β* are fixed, the performance of DANN-DDI is initially improved with the increase of *ɛ* varying from 0 to 0.4 and then drops after that.

**Fig. 2.**
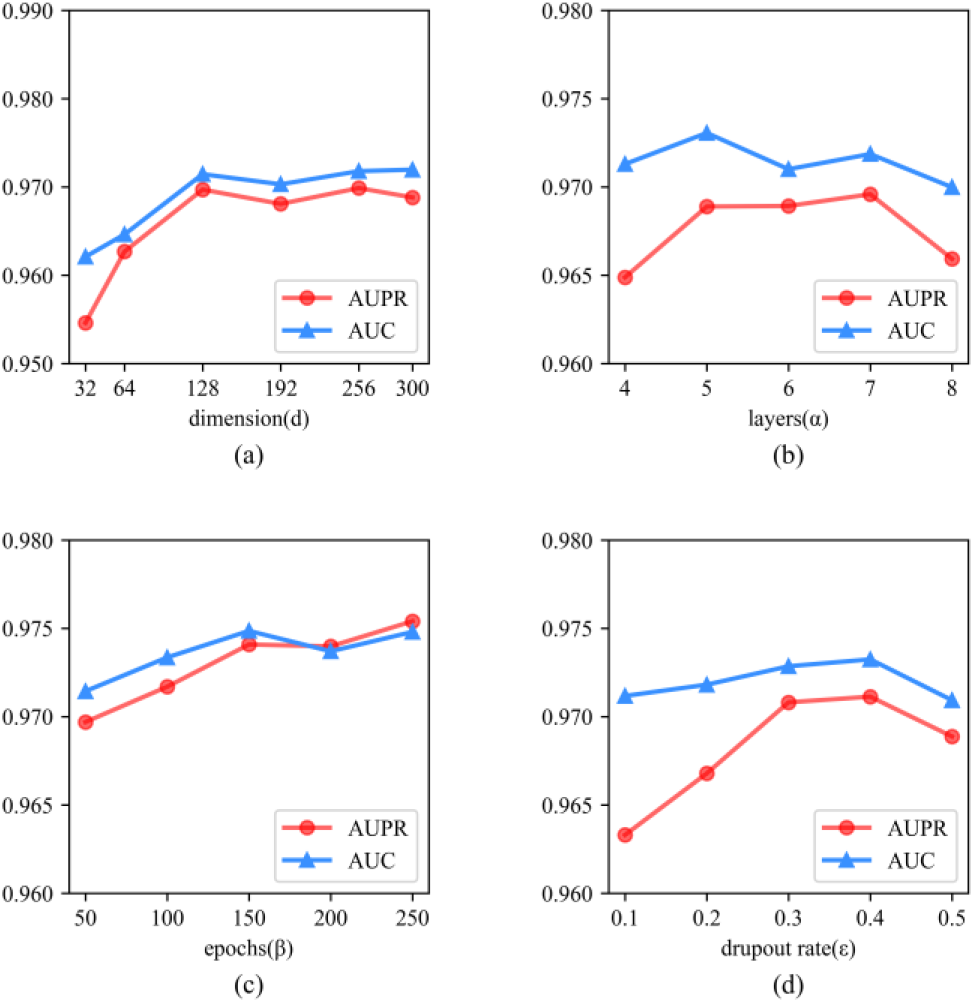
Parameter Sensitivity.

**Fig. 3.**
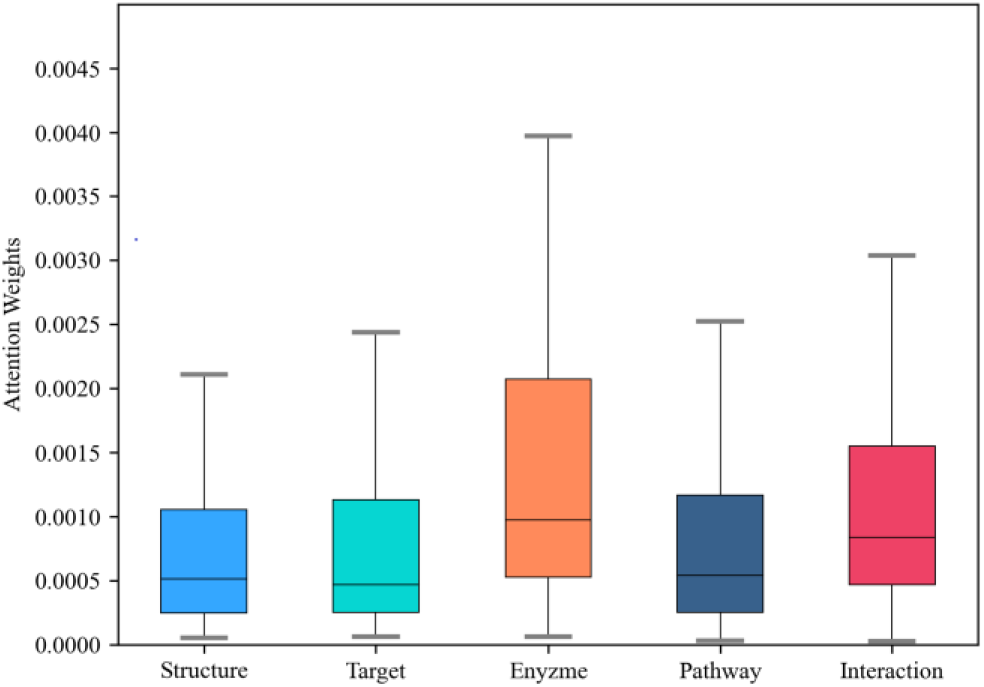
Features’ contributions to the prediction of drug-drug interactions. Substructure, Target, Enzyme, Pathway, and Interaction represent five drug features: chemical substructure, target, enzyme, pathway, and drug-drug interaction respectively.

We adopt *d* = 128, *α* = 7, *β* = 150 and *ɛ* = 0.4 for DANN-DDI in the following studies.

### 3.3 Performances of DANN-DDI

To demonstrate good performances and robustness of DANN-DDI, we investigate three critical components of DANN-DDI: drug feature learning, drug-drug pair feature learning and drug-drug interaction prediction. Here, we attempt to demonstrate that they are the leading factors for the success of DANN-DDI.

#### 3.3.1 Drug Feature Learning

Drug feature learning aims to learn drug embeddings from multiple drug feature networks. Graph embedding methods in machine learning can be adopted in the drug feature learning component. We replace SDNE with several state-of-the-art graph embedding methods: Node2vec [44], Meta-path2vec [45] and LINE [46]. The 5-CV results based on different graph embedding methods are reported in Table 2.

**TABLE 2.** Performances of dann-ddi

| DRL | AUPR | AUC | F | Accuracy | Precision | Recall |
| --- | --- | --- | --- | --- | --- | --- |
| Node2vec+ANN+DNN | 0.9595 | 0.9646 | 0.9565 | 0.9947 | 0.9846 | 0.9301 |
| Metapath2vec+ANN+DNN | 0.9173 | 0.9026 | 0.8934 | 0.9874 | 0.9579 | 0.8371 |
| LINE+ANN+DNN | 0.9639 | 0.9686 | 0.9613 | 0.9952 | 0.9857 | 0.9381 |
| SDNE+ANN+DNN | **0.9709** | **0.9763** | **0.9692** | **0.9962** | **0.9853** | **0.9537** |
| SDNE+DNN | 0.9372 | 0.9462 | 0.9305 | 0.9865 | 0.9680 | 0.8958 |
| SDNE+ANN+DNN | **0.9709** | **0.9763** | **0.9692** | **0.9962** | **0.9853** | **0.9537** |
| SDNE+SVM | 0.4358 | 0.6972 | 0.4171 | 0.9228 | 0.3972 | 0.4390 |
| SDNE+ LR | 0.4425 | 0.7086 | 0.4217 | 0.9193 | 0.3839 | 0.4677 |
| SDNE+ RF | 0.8994 | 0.8940 | 0.8805 | 0.9864 | **0.9973** | 0.7881 |
| SDNE+ DNN | **0.9372** | **0.9462** | **0.9305** | **0.9865** | 0.9680 | **0.8958** |

DRL = Drug representation learning, ANN = Attention neural network, DNN = Deep neural networks.

From the results, we observe that DANN-DDI using SDNE performs better than Node2vec and LINE. The representations generated by the Metapath2vec lead to the worst DDI prediction performance. The results show the effectiveness of SDNE in the DANN-DDI framework.

#### 3.3.2 Drug-drug Pair Feature Learning

Drug-drug pair feature learning aims to learn effective representations for drug-drug pairs. An attention neural network is adopted in this component, which takes into account different contributions of different features and their dimensions. In order to validate the effectiveness of the attention neural network, the comparison method (SDNE+DNN) directly merge the embedding vectors of drugs to represent their pairs.

The performances of the two methods are shown in Table 2. We observe that DANN-DDI achieves a 3.60% improvement in terms of AUPR and a 3.18% improvement in terms of AUC over the comparison method. The results indicate that the attention neural network in DANN-DDI can efficiently learn representations of drug-drug pairs.

The representation of drug-drug pairs has the same dimensions corresponding to the representation of the drugs in pairs. To observe different contributions of the five drug features intuitively, we output the attention weights of all drug pairs from one fold in 5-CV and calculate the average weight of each dimension. The attention weights of 128 dimensions related to each feature are adopted, and we visualize the average weights of all drug pairs on the five features. As shown in Figure 3, we observe that the median attention weight of the enzyme is the highest, followed by the interaction, while target, structure, and pathway have lower median attention weights. In addition, enzyme and interaction features have higher attention weights on the whole. Therefore, enzyme and interaction may have more contributions to the representation of drug-drug pairs and may be more important to the prediction of drug-drug interactions.

#### 3.3.3 Drug-drug Interaction Prediction

The drug-drug interaction prediction component utilizes the representations of drug-drug pairs to predict potential DDIs. In the design of DANN-DDI, we adopt the classic deep neural network (DNN) as the component. In order to test the performance of the deep neural network, we replace DNN with several classic classifiers: Random Forest (RF) [47], Support Vector Machine (SVM) [48] and Logistic Regression classifier (LR) [49]. The learned drug embeddings of the two drugs in drug-drug pairs *E*_*i*_ and *E*_*j*_ are directly concatenated as [*E*_*i*_,*E*_*j*_]. Then we feed the drug-drug pairs’ representation into different classifiers to evaluate the performances.

Table 2 shows the superiority of DNN over the compared classifiers, from which DNN achieves a 4.20% improvement in terms of AUPR and a 5.84% improvement in terms of AUC compared to RF. Overall, conventional classifiers SVM and LR cannot extract high-level drug features, and results in the worst performances. The Random Forest classifier adopts the ensemble learning strategy to handle the non-linear drug feature data. Thus, it performs the second-best in terms of AUPR.

### 3.4 Comparison with DDI Prediction Methods

In this section, we compare the proposed model DANN-DDI with several state-of-the-art prediction methods and baseline methods. We adjust the parameters of these comparison methods to ensure optimal performances.

#### 3.4.1 Compared Methods

First, we compare the following state-of-the-art methods with DANN-DDI to evaluate the performances.

- **Nearest Neighbor**[50] uses the nearest neighbor similarity of drugs contained in known drug-drug interactions to predict new DDIs.
- **Label Propagation**[15] is a semi-supervised learning method based on graph which propagates the existing drug-drug interaction information in the network to identify novel DDIs.
- **MP**[43] is a matrix perturbation method assuming that random removal of a small proportion of links from the network will not alter the network structure.
- **LPMS**[14] is an integrative label propagation method which integrates multiple similarities derived from drug chemical substructures, drug label side effects and off-label side effects.
- **NDLM**[26] uses autoencoders and a deep feed-forward network that are trained using four Jaccard similarity matrixes of four drug features to predict DDIs.
- **DDI-MDAE**[22] utilizes multi-modal deep auto-encoders to learn unified drug representations in a shared hidden layer from multiple drug feature networks. We learn drug representations using the representation learning method in DDI-MDAE and feed them to DNN to predict DDIs.
- **DANN-DDI\*** adopts a single heterogeneous network which integrates multiple drug feature information as the input of representation learning, and the drug-drug pair feature learning and drug-drug interaction prediction components remain unchanged.

In addition, due to the imbalanced dataset, we add simpler baseline methods to provide more comprehensive comparison. In the experiment, each drug can be represented by a binary feature vector, whose values (1 or 0) indicate the presence or absence of the corresponding feature. Then there are two strategies: one is to utilize principal component analysis (PCA) to reduce dimensions; the other is to calculate the pairwise drug-drug similarity using Jaccard similarity measure. Then feature vectors pass through NB, SVM, and LR respectively.

#### 3.4.2 Cross validation results of compared methods

To avoid the bias of results, we implement 10 runs of 5-CV for each model respectively, and adopt the average results of the 10 runs to measure the performances.

As mentioned above, we adopt AUPR as the primary evaluation metric. As shown in Table 3, DANN-DDI outperforms all the comparison methods in most cases. DANN-DDI performs slightly better than NDLM and DDI-MDAE in terms of AUPR and F-measure on 5-CV. In addition, DANN-DDI achieves a 3.45% improvement over MP in terms of AUPR and an 8.63% improvement over MP in terms of F-measure. Simple baseline methods generally report dismal results on our dataset. Moreover, standard deviation for multiple CV runs shows the robustness of DANN-DDI. From the overall results, we observe that our framework DANN-DDI achieves satisfactory performances in DDI prediction.

**TABLE 3.** 5-FOLD CROSS-VALIDATION RESULTS OF DIFFERENT MODELS

| Method | AUPR | AUC | F | Accuracy | Precision | Recall |
| --- | --- | --- | --- | --- | --- | --- |
| PCA+NB | 0.6962+/-0.0002 | 0.3742+/-0.0002 | 0.2142+/-0.0001 | 0.5058+/-0.0004 | 0.3009+/-0.0002 | 0.8648+/-0.0000 |
| PCA+SVM | 0.6994+/-0.0002 | 0.4344+/-0.0004 | 0.3969+/-0.0003 | 0.4395+/-0.0005 | 0.4171+/-0.0004 | 0.9293+/-0.0000 |
| PCA+LR | 0.7112+/-0.0002 | 0.4418+/-0.0003 | 0.3849+/-0.0002 | 0.4680+/-0.0004 | 0.4224+/-0.0003 | 0.9264+/-0.0000 |
| Jaccard+NB | 0.4405+/-0.0000 | 0.7360+/-0.0000 | 0.1541+/-0.0000 | 0.7102+/-0.0007 | 0.2532+/-0.0000 | 0.7589+/-0.0000 |
| Jaccard+SVM | 0.6995+/-0.0001 | 0.4347+/-0.0003 | 0.3975+/-0.0003 | 0.4396+/-0.0003 | 0.4175+/-0.0003 | 0.9294+/-0.0000 |
| Jaccard+LR | 0.7116+/-0.0001 | 0.4428+/-0.0002 | 0.3864+/-0.0001 | 0.4687+/-0.0003 | 0.4236+/-0.0002 | 0.9266+/-0.0000 |
| Nearest Neighbor | 0.6344+/-0.0001 | 0.9381+/-0.0000 | 0.6102+/-0.0003 | 0.9579+/-0.0002 | 0.6601+/-0.0043 | 0.5676+/-0.0032 |
| Label Propagation | 0.6192+/-0.0002 | 0.9348+/-0.0000 | 0.6026+/-0.0003 | 0.9567+/-0.0002 | 0.6449+/-0.0036 | 0.5657+/-0.0027 |
| LPMS | 0.3826+/-0.0003 | 0.8634+/-0.0001 | 0.4166+/-0.0005 | 0.9239+/-0.0011 | 0.3758+/-0.0045 | 0.4681+/-0.0070 |
| MP | 0.9385+/-0.0004 | **0.9854+/-0.0002** | 0.8922+/-0.0005 | 0.9878+/-0.0001 | 0.9180+/-0.0022 | 0.8678+/-0.0019 |
| NDLM | 0.9664+/-0.0011 | 0.9772+/-0.0006 | 0.9650+/-0.0011 | 0.9960+/-0.0001 | 0.9743+/-0.0028 | **0.9559+/-0.0012** |
| DDI-MDAE | 0.9634+/-0.0017 | 0.9625+/-0.0024 | 0.9587+/-0.0021 | 0.9926+/-0.0004 | **0.9945+/-0.0016** | 0.9255+/-0.0048 |
| DANN-DDI* | 0.9415+/-0.0030 | 0.9567+/-0.0013 | 0.9382+/-0.0029 | 0.9924+/-0.0004 | 0.9619+/-0.0070 | 0.9159+/-0.0028 |
| DANN-DDI | **0.9709+/-0.0019** | 0.9763+/-0.0010 | **0.9692+/-0.0019** | **0.9962+/-0.0002** | 0.9853+/-0.0033 | 0.9537+/-0.0020 |

### 3.5 Case Studies

#### 3.5.1 Predicting New Drug-drug interactions

In this section, we conduct experiments to demonstrate the efficacy of DANN-DDI to predict novel DDIs.

In the experiment, the 841 drugs collected have 353,220 drug-drug pairs. Specifically, there are 82,620 known DDIs and 270,600 unlabeled pairs that may contain potential DDIs between drugs. We train DANN-DDI with all known drug-drug interactions to predict DDIs that are not contained in our datasets.

Table 4 demonstrates the top 20 predicted drug-drug interactions by DANN-DDI. We search for the evidence to support the findings, and 14 novel interactions are confirmed in the 5.1.7 version of the DrugBank database (accessed on July 19, 2020). For example, it was reported that Rufinamide may increase the central nervous system depressant (CNS depressant) activities of Fluoxetine, and the studies revealed that the metabolism of Paclitaxel can be decreased when combined with Bromocriptine. The description of the interaction between Fluoxetine (DB00472) and Argatroban (DB00278) is “The risk or severity of hemorrhage can be increased when Fluoxetine is combined with Argatroban.”. The case studies demonstrate the ability and power of DANN-DDI for predicting unobserved drug-drug interactions.

**TABLE 4.**
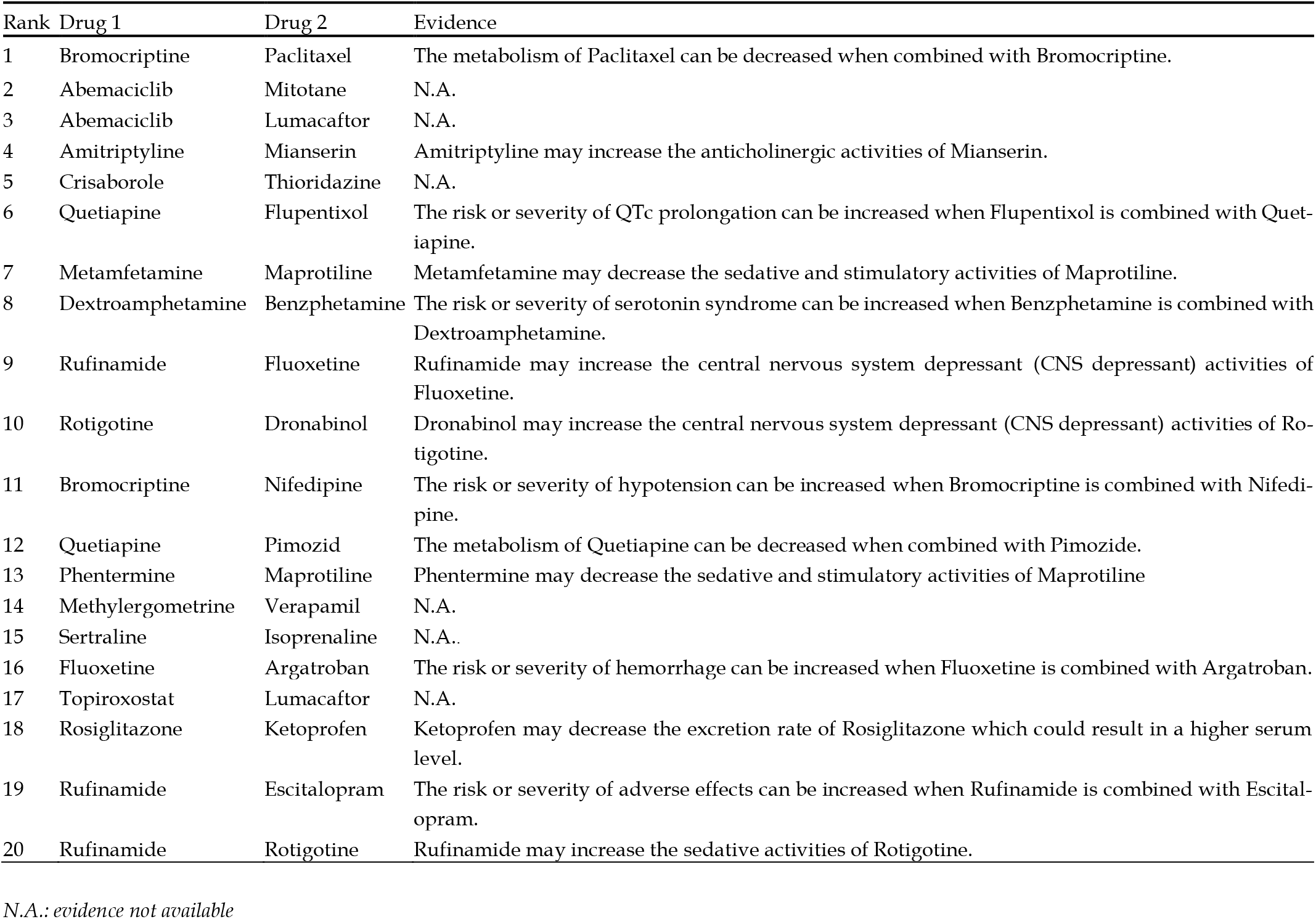
Top 20 ddis predicted by dann-ddi

#### 3.5.2 Predicting New DDI-associated Events

Predicting DDI-associated events is helpful to understand the underlying mechanism of adverse reactions, and to guide the combination of drugs. Hence, we extend DANN-DDI to predict new DDI-associated events. The drug feature learning and drug-drug pair feature learning components of our model for DDI-associated event prediction stay the same as those for binary DDI prediction. For the settings for deep neural network, we set the number of neurons in the output layer as 65, denoting 65 types of events. We select the softmax function to generate probabilities for the output nodes. In addition, cross-entropy loss function is adopted. We utilize the interaction event dataset by [27] and adopt the same evaluation metrics. We apply 5-fold cross-validation (5-CV) to predict DDI-associated events between known drugs. The experimental results show that DANN-DDI can achieve an accuracy score of 0.8874, a micro AUPR score of 0.9088, a micro AUC score of 0.9943 and a macro-F1 score of 0.7781. Performances of DANN-DDI are close to those of [27].

Furthermore, we conduct case studies to validate the effectiveness of our model in practice. We utilize all DDIs and their associated events in [27] to train the prediction model, and then predict associated events for other drug-drug pairs. DDI-associated events and corresponding descriptions are illustrated in [27]. We are concerned with the ten most frequent events, numbered from # 1 to # 10, and examine the top 20 predictions associated with each event. We search for the evidence of the newly predicted DDI-associated events in the latest version 5.1.7 of DrugBank database.

We list ten confirmed drug-drug interaction events in Table 5. For instance, the interaction between Etodolac and Naproxen is predicted to cause event #2. The interaction between the two drugs is described as “The risk or severity of adverse effects can be increased when Etodolac is combined with Naproxen.”. The description of interaction which may lead to event #5 between Homatropine and Donepezil is “The therapeutic efficacy of Homatropine can be decreased when used in combination with Donepezil.”. The case studies demonstrate the promise of our method to detect the potential DDI-associated events.

**TABLE 5.** Confirmed DDIs and their associated events

| Event type | Drug names | Events |
| --- | --- | --- |
| #1 | Nateglinide and Clemastine | The metabolism of Nateglinide can be decreased when combined with Clemastine. |
| #2 | Etodolac and Naproxen | The risk or severity of adverse effects can be increased when Etodolac is combined with Naproxen. |
| #3 | Halofantrine and Mifepristone | The serum concentration of Halofantrine can be increased when it is combined with Mifepristone. |
| #4 | Cyproterone acetate and Bosentan | The serum concentration of Cyproterone acetate can be decreased when it is combined with Bosentan. |
| #5 | Homatropine and Donepezil | The therapeutic efficacy of Homatropine can be decreased when used in combination with Donepezil. |
| #6 | Buprenorphine and Hydromorphone | Hydromorphone may increase the central nervous system depressant (CNS depressant) activities of Buprenorphine. |
| #7 | Amoxapine and Dronedaron | Dronedaron may increase the QTc-prolonging activities of Amoxapine. |
| #8 | Bisoprolol and Diazoxide | Diazoxide may increase the hypotensive activities of Bisoprolol. |
| #9 | Carbamazepine and Dronedaron | The metabolism of Carbamazepine can be increased when combined with Dronedaron. |
| #10 | Bisoprolol and Diclofenac | Diclofenac may decrease the antihypertensive activities of Bisoprolol. |

## 4 Conclusion

In this paper, we study how to attentively integrate multiple drug features to predict unknown drug-drug interactions and then propose a deep attention neural network framework named DANN-DDI. We evaluate the performance of our proposed model against several state-of-the-art prediction methods and baseline methods. Extensive experimental results demonstrate the superiority of DANN-DDI compared to the comparison methods and the efficacy of the attention mechanism. More importantly, the proposed model can also find novel interactions and DDI-associated events. In the future, we will consider optimizing and improving our work from the following aspects: (1) In the original data set, imbalanced data and the noise in the data bring challenges to the prediction accuracy. The future direction is to solve the problem of the imbalanced learning and enhance the robustness to noise; (2) Our method can be used to predict DDI-associated events, but there is still room for improvement. Some events lack detailed descriptions and proved interactions, and the interpretability of the model is needed. Enhancing the accuracy of DDI-associated event prediction can also be our future work.

## ACKNOWLEDGMENT

This work was supported by the Fundamental Research Funds for the Central Universities (2662019QD011), National Key Research and Development Program of China (2018YFC1604000), the National Natural Science Foundation of China (61772381), Huazhong Agricultural University Scientific & Technological Self-innovation Foundation. The funders have no role in study design, data collection, data analysis, data interpretation, or writing of the manuscript.

## APPENDIX

The source codes of DANN-DDI are available at the following GitHub repository. https://github.com/naodandandan/DANN-DDI

